# Cell specificity of adeno-associated virus (AAV) serotypes in human cortical organoids

**DOI:** 10.1101/2023.04.13.536491

**Authors:** Morgan M. Stanton, Harsh N. Hariani, Jordan Sorokin, Patrick M. Taylor, Sara Modan, Brian G. Rash, Sneha B. Rao, Luigi Enriquez, Daphne Quang, Pei-Ken Hsu, Justin Paek, Dorah Owango, Carlos Castrillo, Justin Nicola, Pavan Ramkumar, Andy Lash, Douglas Flanzer, Kevan Shah, Saul Kato, Gaia Skibinski

## Abstract

Human-derived cortical organoids (hCOs) recapitulate cell diversity and 3D structure found in the human brain and offer a promising model for discovery of new gene therapies targeting neurological disorders. Adeno-associated viruses (AAVs) are the most promising vehicles for non-invasive gene delivery to the central nervous system (CNS), but reliable and reproducible *in vitro* models to assess their clinical potential are lacking. hCOs can take on these issues as they are a physiologically relevant model to assess AAV transduction efficiency, cellular tropism, and biodistribution within the tissue parenchyma, all of which could significantly modulate therapeutic efficacy. Here, we examine a variety of naturally occurring AAV serotypes and measure their ability to transduce neurons and glia in hCOs from multiple donors. We demonstrate cell tropism driven by AAV serotype and hCO donor and quantify fractions of neurons and astrocytes transduced with GFP as well as overall hCO health.

## Introduction

Adeno-associated viruses (AAVs) are non-pathogenic viruses packing single-stranded DNA capable of infecting dividing and non-dividing cells. Significant progress has been made in utilizing AAVs for gene therapy delivery in neurological disease as they are capable of gene replacement for loss-of-function mutations and gene silencing for gain-of-function mutations^1–4^. Furthermore, clinical headway of AAVs is evident in the current 27 gene therapy products approved by the Food and Drug Administration (FDA) and a prediction the FDA will approve another 10-20 new gene therapy products every year after 2025^5,6^. AAVs are advantageous for targeting the central nervous system (CNS) and chronic neurological diseases due to their low cytotoxicity, low immunogenicity, and persistent transgene expression in post-mitotic cells^7^. However, gene therapy research with AAVs can often lack translational relevance due to the discrepancy between results obtained in rodent models, non-human primates (NHPs), and humans^8–11^. While considered to be the most translatable animal models, NHP models for gene therapy testing have low experimental throughput, high experimental costs, and raise ethical challenges^12,13^, further motivating the need for human relevant alternative models.

Induced pluripotent stem cell (iPSC) derived hCOs represent a promising complex *in vitro* human model class to study AAV gene therapies as they contain organized three-dimensional tissue architecture and a diversity of CNS cell types, including excitatory neurons, inhibitory neurons, astrocytes, and choroid plexus cells^14–16^. Organoid cell diversity and self-organized tissue structure yield a physiologically realistic system that has been used to model various neurological disease phenotypes including Alzheimer’s, frontotemporal dementia, and schizophrenia^17–19^. hCOs can serve as a useful non-animal model to examine AAVs and provide human physiological insight into AAV transduction efficiency, toxicity, and cell tropism in the CNS^20^. However, there has been limited research to date utilizing brain organoids for AAV experimentation, particularly for human translatability studies^21,22^.

To demonstrate the advantages of hCOs for quantifying AAVs therapeutic effect, we have cultured hCOs from three independent iPSC clones - including healthy and Alzheimer’s donors - and transduced them with six naturally occurring AAV serotypes containing a GFP payload driven by a CAG promoter (AAV1, AAV2, AAV5, AAV6, AAV8, AAV9). Each AAV serotype has specific cell receptors that have previously shown unique cell-type tropism profiles^2,23,24^. For each AAV serotype, organoid health, transduction efficiency, and cell-type tropism were examined and compared across all three hCO clones. We establish serotype correlated cell-tropism in human hCOs and demonstrate transduction results that correlate with results from NHP studies and oppose results from rodent studies. Our human organoid model platform represents a novel scalable approach to human translatability studies for AAV therapeutic discovery and development.

## Results

### Exploration of AAV serotypes with human derived hCOs

hCOs were differentiated from iPSCs using a single-SMAD organoid protocol. Three different lineages of hCOs were generated: a wild type (WT) clone with no associated disease risk factors, an Alzheimer’s associated clone with a presenilin-1 (PSEN1) mutation (AD), and an isogenic clone (Iso) corrected from the AD line. At hCO culture day 109, each clone was incubated with different AAV serotypes (AAV2, AAV4, AAV5, AAV6, AAV8, or AAV9) containing a CAG promoter and GFP payload (Fig. 1a). All hCOs were incubated with 1×10^11^ GC/mL of virus. Viral concentration was determined through an initial viral dosing test that ranged from 1×10^9^ to 1×10^11^ GC/mL with subsequent GFP quantification (Supplemental Fig. 1). Success of AAV transduction varied between hCO clones and serotypes and was quantified via live imaging of GFP fluorescence. AAV6 had the most successful transduction for all three clones at 9 and 42 days post transduction (Fig. 1b and Supplemental Fig 2). The AAV2, AAV8, and AAV9 serotypes transduced two out of the three clones, but AAV1 and AAV5 had sparse GFP labeling or no visible GFP labeled cells in all three clones (Supplemental Fig. 2). Harvest of the hCOs 42 days post transduction for sectioning and immunohistological staining revealed the uniformity and depth of the AAV-GFP penetration into neural tissue (Fig. 1c). To quantify the depth of GFP penetration, a GFP segmentation mask was generated to measure the integration distance of GFP-labeled cells into the core of the hCO section (Fig. 1d); where a lower value indicates high penetrance depth. The integration distance was calculated for AAV2, AAV6, and AAV9 serotypes as they demonstrated the highest levels of GFP observed during live imaging. No significant differences were found between the serotypes, however, AAV9 showed a trend towards greater GFP penetration compared to the other serotypes (Fig. 1e). To monitor CO health and potential cell death after AAV addition, lactate dehydrogenase (LDH) assays were performed on hCO media collected before AAV addition and multiple time points after (Fig. 1f). Only a slight increase in LDH values was observed 3 days post AAV addition, but this was not significant compared to the control group without genetic modification. None of the AAV serotypes induced cell death over the 42 day culture period for any of the three hCO clones.

**Figure 1.**
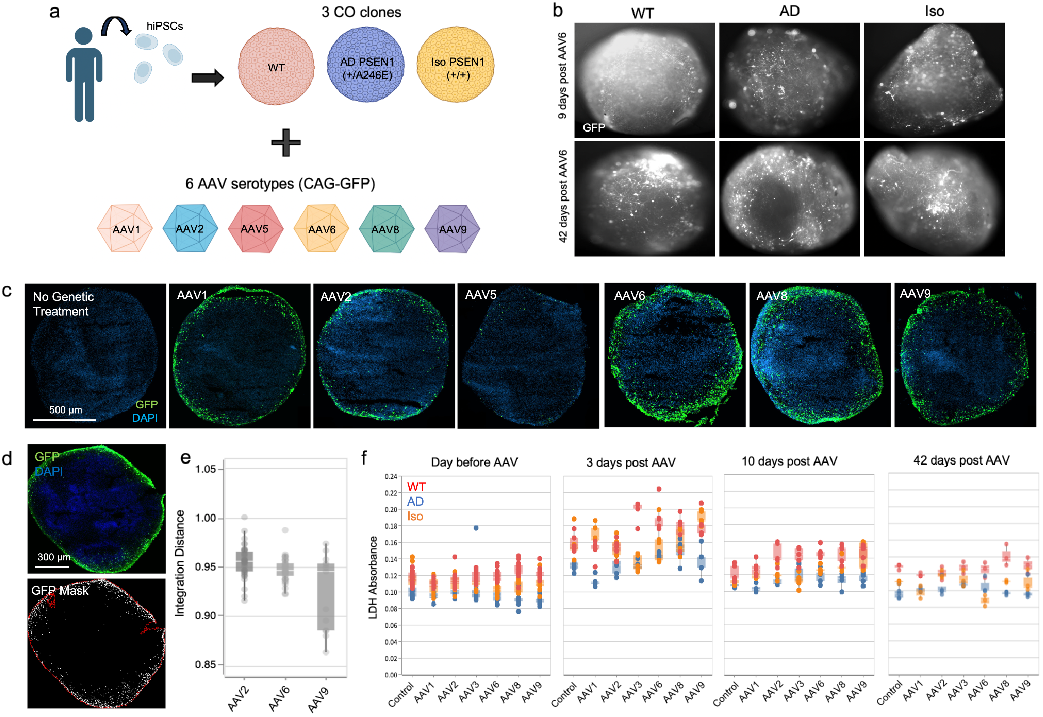
AAV transduction of human cortical organoids. (**a**) hCOs generated from three independent hiPSCs clones (WT, AD, Iso) were transduced with six different AAV serotypes (AAV1, AAV2, AAV5, AAV6, AAV8, AAV9) driven by a CAG promoter and expressing a GFP payload. (**b**) Live hCO GFP imaging 9 and 42 days post AAV6 transduction in hCOs from WT, AD, and Iso hCO clones. hCOs were culture day 118 and 151 respectively. (**c**) Example of GFP transduction 42 days post virus addition of all AAV serotypes in immunohistological stained sections of the Iso hCO clone (culture day 151). Includes a control group without any AAV treatment. (**d**) Example of GFP mask generated from GFP segmentation analysis on a hCO section transduced with AAV6. (**e**) Quantification of Integration distance in hCO sections transduced with AAV2, AAV6, or AAV9. A lower Integration Distance indicates greater penetration of AAV-GFP into the center of the section. (**f**) LDH absorbance measurements before AAV treatment and 3, 10, and 42 days post AAV treatment. No significant differences between parameters (statistical significance determined by two-sided *t*-test).

### Quantification of AAV serotype transduction specificity

To assess which hCO cell types were being transduced, hCOs with each AAV serotype were harvested for immunohistological staining and flow cytometry. Co-staining of GFP and MAP2 (Fig. 2a) or GFP and GFAP (Fig. 2b) of hCO sections revealed transduction of both neurons and astrocytes using the AAV6 serotype. To quantify the transduction efficiencies for all serotypes in all three hCO clones, transduced hCOs were harvested and dissociated for flow cytometry assays. The proportion of GFP-positive cell populations reveal the AAV6 serotype had the greatest transduction efficiency overall compared to the other serotypes (Fig. 2c and Supplemental Fig. 3). However, there was some transduction variability between each clone, with average GFP levels at 6.57%, 10.0%, and 13.1% for WT, AD, and Iso clones respectively. To quantify cell specificity of GFP transduction, cells co-labeled with GFP and MAP2 were measured with flow cytometry to determine the proportion of neurons transduced (Fig. 2d). Within each clone, AAV6, AAV8, and AAV9 had the highest transduction efficiencies of neurons, while AAV5 had little or no GFP-MAP2 labeling. Percentages of neurons transduced varied based on hCO clone, but transduction trends were consistent. Flow cytometry measurements of cells co-labeled with GFP and GFAP were used to quantify the proportion of astrocytes transduced in the hCO (Fig. 2e). Here, there was less variability of efficiency between serotypes and between clones, with only a slight increase in efficiency of AAV6 and AAV8 in the Iso clone.

**Figure 2.**
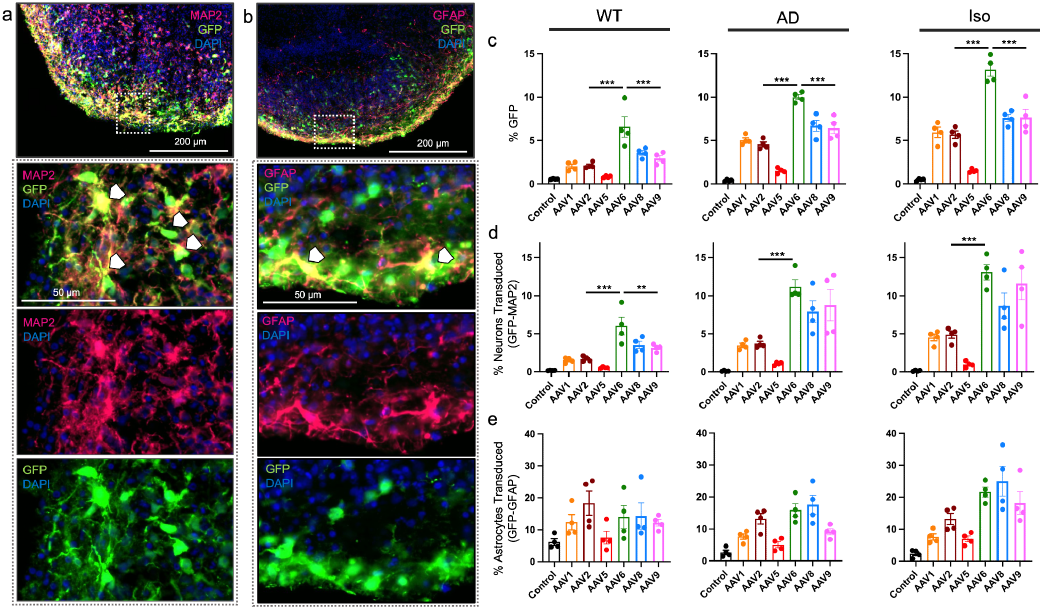
Quantification of AAV cell specificity. (**a & b**) Example of immunohistological sample of Iso clone, 42 days post transduction with AAV6 and labeled with either (a) GFP, MAP2, and DAPI or (b) GFP, GFAP, and DAPI. Dotted square indicates enlarged area. White arrows indicate co-staining of GFP with either (b) MAP2 or (b) GFAP. All hCOs were harvested on day 120, 10 days post AAV transduction. (**c**) Flow cytometry quantification of % GFP labeled cells for each AAV serotype in each hCO clone. (**d**) Flow cytometry quantification of % GFP and MAP2 co-labeled cells for each AAV serotype in each hCO clone. (**e**) Flow cytometry quantification % GFP and GFAP co-labeled cells for each AAV serotype in each hCO clone. All hCOs were harvested on day 120, 10 days post AAV transduction. (*n* = 4 hCOs per condition, statistical significance between AAV6 and AAV2 or AAV9 serotypes determined by one-way ANOVA with Tukey post-hoc tests, \*\*\**p* < 0.001, \*\**p* < 0.01. Values for comparisons between other serotypes are in Supplementary Tables 1-9).

To compare the pairwise performance of each serotype in our hCO model, we used ANOVAs with Tukey’s honestly significant difference (HSD) post hoc tests within each clone for: overall % GFP transduction (Supplementary Tables 1-3), neuronal GFP transduction (%GFP-MAP2 Supplementary Tables 4-6), and astrocyte GFP transduction (%GFP-GFAP, Supplementary Tables 7-9). With regards to transduction efficiency, Using this analysis we observe AAV6 outperforms some of the most commonly used AAVs tested for CNS gene therapies, including AAV2 and AAV9 (Fig 2. c & d).

Summarizing across clones, we used a linear mixed effects model (LMEM) to quantify the effect of AAV6 relative to other serotypes on transduction efficiency of our three measured cell populations (overall % GFP, % GFP-MAP2, and % GFP-GFAP), using serotype as a fixed effect and clone as a random effect. We observed a statistically significant effect of AAV6 relative to all other serotypes for overall % GFP population (Supplementary Table 10, *p* < 0.001 for each comparison) and GFP labeled neurons (% GFP-MAP2) (Supplementary Table 11, *p* < 0.001 for each comparison except AAV9 where *p* < 0.05). In GFP labeled astrocytes (% GFP-GFAP), AAV6 had a statistically significant effect relative to AAV1 and AAV5 (Supplementary Table 12, *p* < 0.001 for each comparison).

## Discussion

### hCO model for examining AAV-based gene therapy

FDA approval rates of compounds targeting the CNS are roughly half of the rates of non-CNS targeting compounds and on average take 30% longer to be approved^25^. Issues for developing CNS AAV therapeutics arise from limited blood-brain-barrier (BBB) penetration and biodistribution across the brain, insufficient understanding of the target and disease mechanisms, and poor *in vitro* models to translate AAV development to humans prior to clinical trials. Human iPSC derived organoids represent the new frontier for personalized medicine and disease modeling as patient samples (healthy or diseased) can be collected, differentiated, and matured into any brain region and tested for therapeutic efficacy and toxicity. They can provide a much needed approach for characterizing AAV therapeutics targeting the CNS, as relevant viable human brain tissue samples are difficult to acquire, maintain in culture, and scale. Early results from Zhou *et al*. validate successful compound screening on forebrain organoids by discovery and validation of novel ZIKA antivirals^26^. However, large-scale screening of AAVs for novel capsid design or therapeutic efficacy with brain organoids has not been documented. The research presented here provides proof-of-concept for utilizing hCOs to measure biodistribution, cell tropism, and cell toxicity of novel AAV therapeutics at scale. We analyzed comparative transduction properties of AAV serotypes. Although some transduction variability between individual hCOs was observed, overall comparative transduction efficiency remained consistent across donors, indicating donor-to-donor reproducibility is not a significant issue. We compared AAV transduction of hCOs with an Alzheimer’s disease related mutation and corresponding isogenic control and found that transduction efficiency of AAV6 was increased in the Iso clone compared to the AD model (Fig. 2). This type of comparison can determine how disease pathogenesis impacts AAV transduction and improve capsid design for strategic therapeutic delivery. Organoid models are continually improving, and ongoing work to advance reproducibility of tissue, increase automation of tissue culture, and quantify the predictivity of organoids for toxicology translation to clinical trials has already enabled them to be applied usefully to therapeutics discovery and development^14,27^.

### AAV transduction of hCOs compared to animal models

In 2022, due to the historical challenges of translatability from animals to humans, the US FDA announced that new drugs can be approved for clinical trials without animal testing,^28^ signaling a new era for therapeutic discovery and validation using human *in vitro* models. Animal models do not capture many aspects of human disease at the tissue, cellular and molecular level and compounds tested for efficacy and/or toxicity in animals often fail in clinical trials^29^. AAV gene therapy has been tested across a range of species including mice and NHPs, such as marmosets, and macaques, but efficacy trials have shown limited reproducibility across species. For example, two generations of BBB-penetrant AAV variants (AAV-PHP.B and AAV-PHP.eB) successfully transduced mice brains but failed in marmosets^8,9^. Moreover, while primate models are typically more physiologically relevant, there are conflicting results between different species of NHPs^11,30^, suggesting the need for a more reliably predictive human experimental model for preclinical development. Complex hCOs provide an alternative, scalable, physiologically relevant human model for studying AAVs.

Using hCOs, our most salient discovery was the success of the AAV6 serotype for overall GFP transduction and transduction of neurons. Eight AAV serotypes have been identified in vertebrates, with AAV2 being the most widely studied serotype for gene therapies targeting the CNS, due to previously documented strong neural tropism. However, very few gene therapies are in clinical trials utilizing AAV6^1,7^. In all hCO donors, the AAV6 serotype outperformed the AAV2 serotype in transducing neurons by nearly two-fold and demonstrated statistical significance within and across clones in our cell populations (Supplementary Tables 1-12). Previous studies have observed successful transduction with AAV6 of motor neurons in African green monkeys^31^ or dorsal root ganglion neurons in marmosets^32^, but systematic comparison of AAV serotypes in mouse models found minimal cell transduction in the CNS with AAV6 and strong neuron labeling with AAV5^33^. In the hCO model, AAV5 was one of the lowest performing serotypes for overall GFP transduction across all clones, illustrating the considerable species-specific differences of AAV transduction success. AAV9 is another commonly applied serotype for AAV gene delivery as it has demonstrated ability to cross the BBB^34^ and is the serotype used with the FDA approved onasemnogene abeparvovec (trade name Zolgensma) for treatment of spinal muscular atrophy. Our hCOs had positive results using AAV9 to transduce neurons as well as astrocytes with GFP. This is similar to previous reports that found macaques injected with AAV9 with GFP had efficient transduction of motor neurons and glial cells across the parenchyma^35^ emphasizing hCO’s ability to recapitulate results in NHPs.

Here, our studies demonstrate the value of hCOs for AAV therapeutic design and development. hCOs are scalable, providing a high throughput screening platform for rapid iterations on AAV design without using animal models. hCOs contain the relevant cell types of the brain, enabling validation of cell type specific transduction and biodistribution. Finally, hCOs can be derived from human patients to identify patient-specific differences in AAV cell type tropism, enabling the use of this system for personalized medicine.

## Materials and Methods

### Cell lines used for generating hCOs

For WT hCO differentiation, a control donor iPSC clone obtained from CIRM (CW20109) that had no history of neuropsychiatric disease and no known disease-associated gene variants was used. For the AD and respective Iso hCO differentiation, a donor with PSEN1 A246E mutation and clinical diagnosis of Alzheimer’s was obtained from Coriell (AG25367). Genetic engineering was used to make an isogenic clone (Iso) by correcting the A246E mutation using CRISPR-Cas9 (crRNA sequence: ACCTCCCTGAATGGACTGAG, ssODN sequence:

TTCCTGTGACAAACAAATTATCAGTCTTGGGTTTTACCATATACTGAAATCACAGCCA AGATGAGCCACGCAGTCCATTCAGGGAGGTACTTGATAAACACCAGGGCCATGAGGGCACTA).

### Culture of hCOs

hCOs were grown in individual wells using a novel dorsal forebrain protocol modified from Velasco *et al*.^16^ to improve cortical fate, reliability, and embryoid body survival in round bottom plates. Briefly, iPSCs were thawed concurrently and passaged twice before seeding. Prior to seeding, iPSCs were dissociated when 50–80% confluent and plated in U-bottom 96-well plates at 9×10^3^ cells per well with ROCK inhibitor. hCOs were cultured in neural induction media containing inhibitory molecules added to direct cortical fate via single-SMAD, Wnt signaling inhibition, as well as Shh inhibition. hCOs were fed on automated Biotek feeders and washers. On the fifth week of culture, organoids were switched to a DMEM-based maturation media with the addition of Growth Factor Reduced (GFR) Basement Membrane Matrix Matrigel. On the eighth week of culture, hCOs were fed maturation medium containing BDNF and GDNF with GFR Matrigel. Matrigel was omitted during the ninth week of culture. hCOs were maintained with automated feeding and imaging until harvest.

### AAV transduction of hCOs

All AAV-CAG-GFP serotypes (AAV1, AAV2, AAV5, AAV6, AAV8, and AAV9) were purchased from Vector Biolabs (7071, 7072, 7073, 7074, 7075, and 7076 respectively). For virus addition to day 109 hCOs, all media was removed from organoid wells and replaced with 100 uL media containing 1×10^11^ GC of virus/mL unless otherwise stated. hCOs were incubated in virus-containing media overnight at 37 °C. After 24 hours, 100 μL of fresh culture media was added to each well. For the following feeds, organoids had 65% media changes every day after virus addition with fresh media until hCOs were harvested for analysis.

### Live immunofluorescence imaging

Live imaging of GFP immunofluorescence in hCOs was obtained using an IN Cell Analyzer 2200 (GE Healthcare) at 4x or 10x magnification.

### Lactate Dehydrogenase (LDH) Assay

Media samples were collected from individual hCO wells and run using the Cytoscan LDH Cytotoxicity Assay (G-Biosciences, 786-210) per manufacturer’s instructions. Briefly, 50 μL of hCO media was transferred to single wells of 96-well black, clear-bottom plates (Corning, 3631) followed by 50 μL of G-Biosciences Reaction Mix. Plate was covered and incubated at room temperature for 30 minutes followed by addition of 50 μL of G-Biosciences Stop Solution to each well. Absorbance was immediately read on a Perkin Elmer 2104 EnVision microplate reader.

### Fixed immunofluorescence sectioning, staining and imaging for hCO sections

hCOs were transferred and rinsed with PBS and fixed in 4% paraformaldehyde (Electron Microscopy Sciences, 15710) for 4 hours at 4 °C. After fixation, organoids were rinsed 3 times with PBS, and incubated with 30% w/v sucrose solution overnight with gentle agitation at 4 °C. Samples were embedded into Optimal Cutting Temperature solution (Tissue Tek, 4583), flash frozen on dry ice, and stored at -80 °C until sectioning. Embedded blocks were sectioned into 25 μm slices using an Epredia NX-70 Cryostar Cryostat (Thermo Fisher). For staining, slides were rinsed with PBS, then incubated in 0.25% Triton X-100 with 4% Donkey serum (EMD Millipore, S30) for 1 hour, followed by incubation in primary antibodies overnight: chicken GFP (1:3000, Aves Labs, GFP-1010) and guinea pig MAP2 (1:1000, Millipore-Sigma, MAB3418). After rinsing 5 times with PBS, slides were incubated in secondary antibodies, donkey anti-chicken 647 (1:1000, Thermo Scientific, A78952), donkey anti-guinea pig 555 (1:1000, Biotium, 20276), and DAPI (1:1000, Biovision, B1098-5) for 2 hours at room temperature and rinsed again 5 times with PBS. Slides were then mounted and imaged using Zeiss Axio Scan Z1 with 20x / 0.8 objective and stitched together by the Zeiss Zen software.

### Penetrance analysis of viral fluorescence into the organoid

hCO sections fixed and labeled with GFP and DAPI were used for analysis of GFP penetration depth. Number of GFP objects per section, GFP coverage of the section, and the average fraction of GFP overlap was computed. Distribution statistics including average “Integration Distance” is defined as:

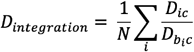

where *D*_*ic*_ is the euclidean distance between GFP puncta *i* and the slice center *c*, and *D*_*bic*_ is the distance between the center *c* and the border pixel *bi* closest to puncta *i*. An Integration Distance of 0.5 means the cell’s position is 1/2 the radius of an idealized circular slice. The lower the value, the more the AAV has penetrated (or cells have migrated) on average.

### Flow cytometry

hCOs for flow cytometry had media removed and washed once with PBS. hCOs were dissociated into single cells using a 1:20 mixture of DNAse in PBS (Worthington Biochemical, LK003163) and TrypLE (Thermo Fisher, 12604013) respectively at 37 °C for 20 minutes followed by gentle manual dissociation with a pipette. An equal volume of 10% v/v fetal bovine serum (FBS, VWR, 45001-108) in PBS was added to the solution to quench the reaction. Cells were centrifuged at 300 rpm and washed 1 time with PBS, placed on ice, and labeled with Live-or-dye 750/777 stain (Biotium, 32008, 1:1000) in cold PBS for 15 minutes. Cells were centrifuged and washed with PBS. Cells were fixed in 2% paraformaldehyde (Electron Microscopy Sciences, 15710) in cold PBS for 20 minutes at 4 °C followed by centrifugation at 1000 rpm, filtration through a 40 μm filter, and washed in PBS. Cells were centrifuged and resuspended in flow buffer for intracellular staining (buffer stock diluted in deionized water (Biolegend, 421002-BL) supplemented with 0.3% v/v BSA (Sigma-Aldrich, A9576), 1% v/v donkey serum (Sigma-Aldrich, S30-100ML) and 0.01% w/v sodium azide. Cells were centrifuged at 1000 rpm and resuspended in flow buffer containing the primary antibodies: rat GFP (Biolegend, 338001, 1:200), guinea pig MAP2 (Synaptic Systems, 188004, 1:4000), and or mouse GFAP-PE (Miltenyi Biotech, 130-118-351, 1:50). Cells were incubated in antibody solution overnight at 4 °C or 30 minutes only for conjugated antibodies. Cells were centrifuged and washed 2 times in cold flow buffer and resuspended in donkey anti-rat 555 secondary antibody (ThermoFisher, A48270, 1:1000), donkey anti-guinea pig 647 secondary antibody (Biotium, 20837, 1:1000), and DAPI (Biovision, B1098-5, 1:10,000) in flow buffer. Cells were incubated for 30 minutes at room temperature, washed 2 times with flow buffer, resuspended in PBS, and stored at 4 °C until flow analysis. Flow cytometry was run on a ThermoFisher Attune NxT with auto sampler and Attune NxT v3.1.2 software. FlowJo v10.6.1 Software was used for data analysis.

### Statistical analysis

For serotype comparisons of flow cytometry data within clones, ANOVAs with Tukey HSD post-hoc tests were performed in R. For serotype comparisons across clones, linear mixed effects models were employed. Here, the iPSC clone is a potential confounding variable and treated as a random effect, while serotype is our fixed effect predictor variable. Mixed effects models were performed using the lmerTest package in R.

## Author Contributions

MM Stanton, HN Hariani, G Skibinski, K Shah, and S Kato conceived and designed experiments. MM Stanton and HN Hariani performed laboratory experiments. D Quang assisted with immunohistological fixing and staining. D Owango, C Castrillo, and J Nicola ran laboratory organoid production pipeline. J Sorokin and P Ramkumar built machine learning models. J Sorokin, P Ramkumar, HN Hariani, K Shah, and MM Stanton ran the analysis. P-K Hsu and J Paek generated isogenic lines. A Lash and D Flanzer built Herophilus laboratory software pipeline. C Castrillo, J Nicola, and D Flanzer built Herophilus automation imaging and feeding infrastructure. MM Stanton and HN Hariani wrote the manuscript with input from G Skibinski, K Shah, and S Kato. All authors discussed the results and reviewed the manuscript.

