## Supplementary Information for "Cell specificity of adeno-associated virus (AAV) serotypes in human cortical organoids"

### Supplemental Information

#### Supplementary Figures

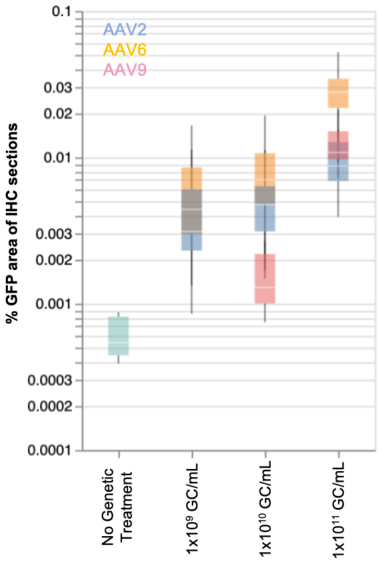


**Supplementary Figure 1.** Dose dependent increase in % GFP positive cell area quantified from immunohistological WT hCO sections comparing AAV2, AAV6 and AAV9 serotypes with CAG-GFP payload (*n* = 4 organoids per condition). hCOs were harvested on day 94, 10 days post AAV transduction. Box plot = upper and lower quartiles, with median center line, error bars = max and min of data set.

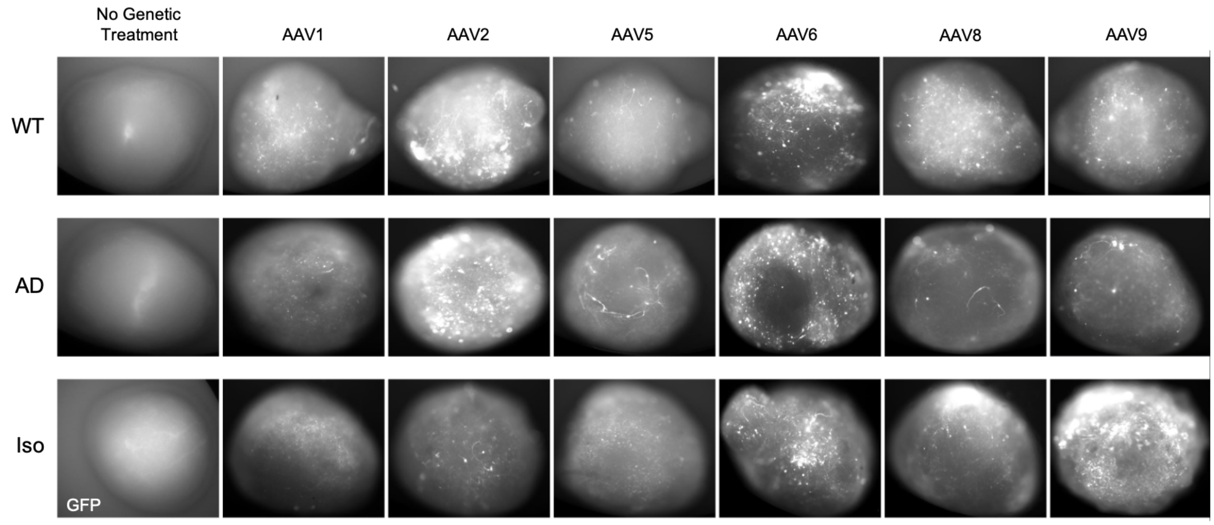


**Supplemental Figure 2.** Live imaging of GFP fluorescence on WT, AD, and Iso hCOs 42 days post transduction with each AAV serotype. hCOs are 151 days old.

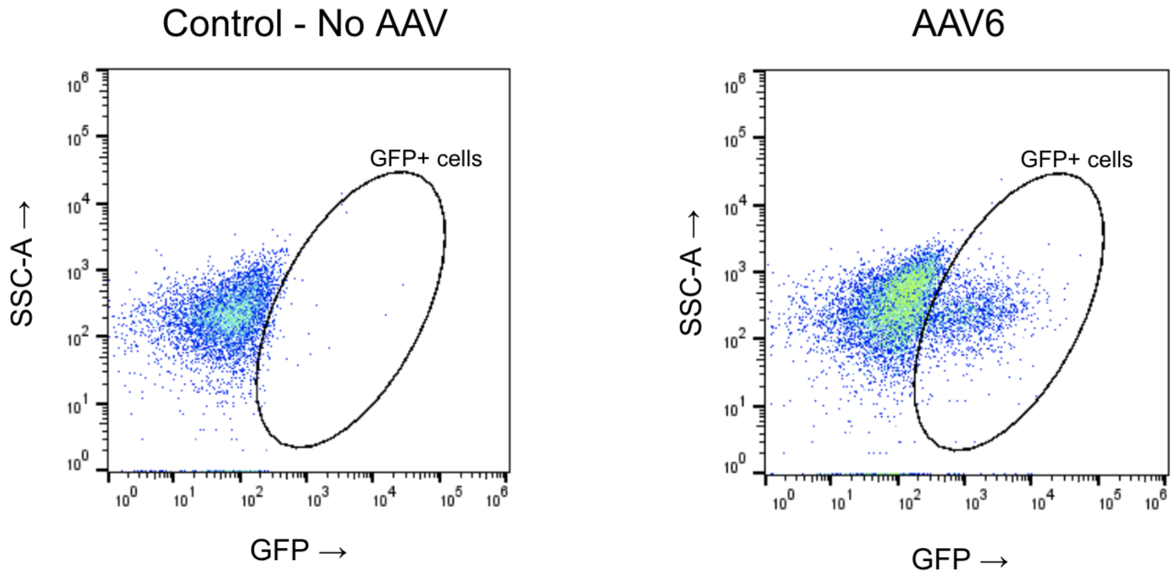


**Supplemental Figure 3.** Example of flow cytometry gating of GFP positive cells from unmodified hCOs (Control) and AAV6 transduced hCOs. hCOs were harvested on day 120, 10 days post AAV transduction.

### Supplemental Tables

### Supplementary Table 1: WT Clone - % GFP Tukey multiple comparisons of means

| Groups compared | Difference between means | Lower bound of confidence interval | Upper bound of confidence interval | adjusted *p*-value |
|---|---|---|---|---|
| *Control-AAV9* | -2.3875 | -4.6368763 | -0.1381237 | 0.0328441 |
| *AAV1-AAV9* | -0.9075 | -3.1568763 | 1.3418763 | 0.8391449 |
| *AAV2-AAV9* | -0.815 | -3.0643763 | 1.4343763 | 0.8947657 |
| *AAV5-AAV9* | -2.1025 | -4.3518763 | 0.1468763 | 0.0769952 |
| *AAV6-AAV9* | 3.6325 | 1.3831237 | 5.8818763 | 0.0005689 |
| *AAV8-AAV9* | 0.64 | -1.6093763 | 2.8893763 | 0.9641365 |
| *AAV1-Control* | 1.48 | -0.7693763 | 3.7293763 | 0.3674218 |
| *AAV2-Control* | 1.5725 | -0.6768763 | 3.8218763 | 0.3018196 |
| *AAV5-Control* | 0.285 | -1.9643763 | 2.5343763 | 0.999527 |
| *AAV6-Control* | 6.02 | 3.7706237 | 8.2693763 | 0.0000004 |
| *AAV8-Control* | 3.0275 | 0.7781237 | 5.2768763 | 0.0041995 |
| *AAV2-AAV1* | 0.0925 | -2.1568763 | 2.3418763 | 0.9999994 |
| *AAV5-AAV1* | -1.195 | -3.4443763 | 1.0543763 | 0.6068447 |
| *AAV6-AAV1* | 4.54 | 2.2906237 | 6.7893763 | 0.0000306 |
| *AAV8-AAV1* | 1.5475 | -0.7018763 | 3.7968763 | 0.3187675 |
| *AAV5-AAV2* | -1.2875 | -3.5368763 | 0.9618763 | 0.5252964 |
| *AAV6-AAV2* | 4.4475 | 2.1981237 | 6.6968763 | 0.0000409 |
| *AAV8-AAV2* | 1.455 | -0.7943763 | 3.7043763 | 0.3864582 |
| *AAV6-AAV5* | 5.735 | 3.4856237 | 7.9843763 | 0.0000009 |
| *AAV8-AAV5* | 2.7425 | 0.4931237 | 4.9918763 | 0.010661 |
| *AAV8-AAV6* | -2.9925 | -5.2418763 | -0.7431237 | 0.0047122 |

**Supplementary Table 2: AD Clone - % GFP Tukey multiple comparisons of means**

| Groups compared | Difference between means | Lower bound of confidence interval | Upper bound of confidence interval | adjusted *p*-value |
|---|---|---|---|---|
| *Control-AAV9* | -6.02 | -7.7905492 | -4.2494508 | 0 |
| *AAV1-AAV9* | -1.385 | -3.1555492 | 0.38554923 | 0.1943067 |
| *AAV2-AAV9* | -1.8625 | -3.6330492 | -0.0919508 | 0.0350712 |
| *AAV5-AAV9* | -4.9125 | -6.6830492 | -3.1419508 | 0.0000002 |
| *AAV6-AAV9* | 3.575 | 1.8044508 | 5.34554923 | 0.0000304 |
| *AAV8-AAV9* | 0.265 | -1.5055492 | 2.03554923 | 0.9987821 |
| *AAV1-Control* | 4.635 | 2.8644508 | 6.40554923 | 0.0000006 |
| *AAV2-Control* | 4.1575 | 2.3869508 | 5.92804923 | 0.0000032 |
| *AAV5-Control* | 1.1075 | -0.6630492 | 2.87804923 | 0.4243986 |
| *AAV6-Control* | 9.595 | 7.8244508 | 11.3655492 | 0 |
| *AAV8-Control* | 6.285 | 4.5144508 | 8.05554923 | 0 |
| *AAV2-AAV1* | -0.4775 | -2.2480492 | 1.29304923 | 0.9722788 |
| *AAV5-AAV1* | -3.5275 | -5.2980492 | -1.7569508 | 0.0000368 |
| *AAV6-AAV1* | 4.96 | 3.1894508 | 6.73054923 | 0.0000002 |
| *AAV8-AAV1* | 1.65 | -0.1205492 | 3.42054923 | 0.0783969 |
| *AAV5-AAV2* | -3.05 | -4.8205492 | -1.2794508 | 0.0002571 |
| *AAV6-AAV2* | 5.4375 | 3.6669508 | 7.20804923 | 0 |
| *AAV8-AAV2* | 2.1275 | 0.3569508 | 3.89804923 | 0.0121174 |
| *AAV6-AAV5* | 8.4875 | 6.7169508 | 10.2580492 | 0 |
| *AAV8-AAV5* | 5.1775 | 3.4069508 | 6.94804923 | 0.0000001 |
| *AAV8-AAV6* | -3.31 | -5.0805492 | -1.5394508 | 0.0000884 |

**Supplementary Table 3: Iso Clone - % GFP Tukey multiple comparisons of means**

| Groups compared | Difference between means | Lower bound of confidence interval | Upper bound of confidence interval | adjusted *p*-value |
|---|---|---|---|---|
| *Control-AAV9* | -7.1625 | -9.6789166 | -4.6460834 | 0.0000001 |
| *AAV1-AAV9* | -1.725 | -4.2414166 | 0.7914166 | 0.3226143 |
| *AAV2-AAV9* | -1.98 | -4.4964166 | 0.5364166 | 0.1893563 |
| *AAV5-AAV9* | -6.14 | -8.6564166 | -3.6235834 | 0.0000018 |
| *AAV6-AAV9* | 5.5025 | 2.9860834 | 8.0189166 | 0.0000095 |
| *AAV8-AAV9* | -0.0475 | -2.5639166 | 2.4689166 | 1 |
| *AAV1-Control* | 5.4375 | 2.9210834 | 7.9539166 | 0.0000114 |
| *AAV2-Control* | 5.1825 | 2.6660834 | 7.6989166 | 0.0000229 |
| *AAV5-Control* | 1.0225 | -1.4939166 | 3.5389166 | 0.8347803 |
| *AAV6-Control* | 12.665 | 10.1485834 | 15.1814166 | 0 |
| *AAV8-Control* | 7.115 | 4.5985834 | 9.6314166 | 0.0000002 |
| *AAV2-AAV1* | -0.255 | -2.7714166 | 2.2614166 | 0.9998699 |
| *AAV5-AAV1* | -4.415 | -6.9314166 | -1.8985834 | 0.0002036 |
| *AAV6-AAV1* | 7.2275 | 4.7110834 | 9.7439166 | 0.0000001 |
| *AAV8-AAV1* | 1.6775 | -0.8389166 | 4.1939166 | 0.3529558 |
| *AAV5-AAV2* | -4.16 | -6.6764166 | -1.6435834 | 0.0004287 |
| *AAV6-AAV2* | 7.4825 | 4.9660834 | 9.9989166 | 0.0000001 |
| *AAV8-AAV2* | 1.9325 | -0.5839166 | 4.4489166 | 0.2103587 |
| *AAV6-AAV5* | 11.6425 | 9.1260834 | 14.1589166 | 0 |
| *AAV8-AAV5* | 6.0925 | 3.5760834 | 8.6089166 | 0.000002 |
| *AAV8-AAV6* | -5.55 | -8.0664166 | -3.0335834 | 0.0000084 |

**Supplementary Table 4: WT Clone - % GFP-MAP2 Tukey multiple comparisons of means**

| Groups compared | Difference between means | Lower bound of confidence interval | Upper bound of confidence interval | adjusted *p*-value |
|---|---|---|---|---|
| *AAV1-Control* | 1.3416177 | -0.9070895 | 3.5903249 | 0.4783297 |
| *AAV2-Control* | 1.5183692 | -0.730338 | 3.7670764 | 0.3389338 |
| *AAV5-Control* | 0.3829972 | -1.8657101 | 2.6317044 | 0.9974978 |
| *AAV6-Control* | 5.8528369 | 3.6041297 | 8.1015442 | 0.0000006 |
| *AAV8-Control* | 3.3764734 | 1.1277661 | 5.6251806 | 0.0013207 |
| *AAV9-Control* | 3.0008558 | 0.7521485 | 5.249563 | 0.004571 |
| *AAV2-AAV1* | 0.1767515 | -2.0719557 | 2.4254587 | 0.9999707 |
| *AAV5-AAV1* | -0.9586205 | -3.2073278 | 1.2900867 | 0.8030932 |
| *AAV6-AAV1* | 4.5112193 | 2.262512 | 6.7599265 | 0.0000334 |
| *AAV8-AAV1* | 2.0348557 | -0.2138516 | 4.2835629 | 0.0931891 |
| *AAV9-AAV1* | 1.6592381 | -0.5894692 | 3.9079453 | 0.2474309 |
| *AAV5-AAV2* | -1.135372 | -3.3840793 | 1.1133352 | 0.6590095 |
| *AAV6-AAV2* | 4.3344677 | 2.0857605 | 6.583175 | 0.0000583 |
| *AAV8-AAV2* | 1.8581042 | -0.3906031 | 4.1068114 | 0.1507953 |
| *AAV9-AAV2* | 1.4824866 | -0.7662207 | 3.7311938 | 0.3652271 |
| *AAV6-AAV5* | 5.4698398 | 3.2211325 | 7.718547 | 0.0000019 |
| *AAV8-AAV5* | 2.9934762 | 0.744769 | 5.2421834 | 0.0046833 |
| *AAV9-AAV5* | 2.6178586 | 0.3691514 | 4.8665658 | 0.0158793 |
| *AAV8-AAV6* | -2.4763636 | -4.7250708 | -0.2276564 | 0.0248558 |
| *AAV9-AAV6* | -2.8519812 | -5.1006884 | -0.6032739 | 0.0074478 |
| *AAV9-AAV8* | -0.3756176 | -2.6243248 | 1.8730896 | 0.9977528 |

**Supplementary Table 5: AD Clone - % GFP-MAP2 Tukey multiple comparisons of means**

| Groups compared | Difference between means | Lower bound of confidence interval | Upper bound of confidence interval | adjusted *p*-value |
|---|---|---|---|---|
| *AAV1-Control* | 3.3882606 | -1.3159627 | 8.092484 | 0.2711967 |
| *AAV2-Control* | 3.6037084 | -1.1005149 | 8.307932 | 0.2125667 |
| *AAV5-Control* | 0.9827411 | -3.7214821 | 5.686964 | 0.9924481 |
| *AAV6-Control* | 11.0771844 | 6.3729611 | 15.781408 | 0.0000031 |
| *AAV8-Control* | 7.8262757 | 3.1220524 | 12.530499 | 0.0003967 |
| *AAV9-Control* | 8.6664191 | 3.9621959 | 13.370642 | 0.0001075 |
| *AAV2-AAV1* | 0.2154478 | -4.4887755 | 4.919671 | 0.9999988 |
| *AAV5-AAV1* | -2.4055195 | -7.1097427 | 2.298704 | 0.6463153 |
| *AAV6-AAV1* | 7.6889238 | 2.9847005 | 12.393147 | 0.0004922 |
| *AAV8-AAV1* | 4.4380151 | -0.2662082 | 9.142238 | 0.0727625 |
| *AAV9-AAV1* | 5.2781585 | 0.5739353 | 9.982382 | 0.0214602 |
| *AAV5-AAV2* | -2.6209672 | -7.3251905 | 2.083256 | 0.5553434 |
| *AAV6-AAV2* | 7.473476 | 2.7692528 | 12.177699 | 0.000691 |
| *AAV8-AAV2* | 4.2225673 | -0.481656 | 8.926791 | 0.0975785 |
| *AAV9-AAV2* | 5.0627108 | 0.3584875 | 9.766934 | 0.0296284 |
| *AAV6-AAV5* | 10.0944432 | 5.39022 | 14.798667 | 0.0000126 |
| *AAV8-AAV5* | 6.8435345 | 2.1393113 | 11.547758 | 0.0018704 |
| *AAV9-AAV5* | 7.683678 | 2.9794547 | 12.387901 | 0.0004962 |
| *AAV8-AAV6* | -3.2509087 | -7.955132 | 1.453315 | 0.3139909 |
| *AAV9-AAV6* | -2.4107653 | -7.1149885 | 2.293458 | 0.6441153 |
| *AAV9-AAV8* | 0.8401435 | -3.8640798 | 5.544367 | 0.9967528 |

**Supplementary Table 6: Iso Clone - % GFP-MAP2 Tukey multiple comparisons of means**

| Groups compared | Difference between means | Lower bound of confidence interval | Upper bound of confidence interval | adjusted *p*-value |
|---|---|---|---|---|
| *AAV1-Control* | 4.3885621 | -0.7543279 | 9.5314521 | 0.1282969 |
| *AAV2-Control* | 4.759172 | -0.383718 | 9.9020621 | 0.0817652 |
| *AAV5-Control* | 0.8543156 | -4.2885744 | 5.9972056 | 0.9978206 |
| *AAV6-Control* | 12.9633571 | 7.8204671 | 18.1062471 | 0.000001 |
| *AAV8-Control* | 8.5401709 | 3.3972809 | 13.6830609 | 0.0004058 |
| *AAV9-Control* | 11.4834874 | 6.3405974 | 16.6263774 | 0.000007 |
| *AAV2-AAV1* | 0.3706099 | -4.7722801 | 5.5134999 | 0.9999824 |
| *AAV5-AAV1* | -3.5342465 | -8.6771365 | 1.6086435 | 0.3199454 |
| *AAV6-AAV1* | 8.574795 | 3.431905 | 13.717685 | 0.0003862 |
| *AAV8-AAV1* | 4.1516087 | -0.9912813 | 9.2944987 | 0.1685259 |
| *AAV9-AAV1* | 7.0949253 | 1.9520353 | 12.2378153 | 0.0032723 |
| *AAV5-AAV2* | -3.9048565 | -9.0477465 | 1.2380335 | 0.2205995 |
| *AAV6-AAV2* | 8.204185 | 3.061295 | 13.347075 | 0.0006581 |
| *AAV8-AAV2* | 3.7809988 | -1.3618912 | 8.9238888 | 0.2509411 |
| *AAV9-AAV2* | 6.7243154 | 1.5814254 | 11.8672054 | 0.0055799 |
| *AAV6-AAV5* | 12.1090415 | 6.9661515 | 17.2519315 | 0.0000031 |
| *AAV8-AAV5* | 7.6858553 | 2.5429653 | 12.8287453 | 0.0013919 |
| *AAV9-AAV5* | 10.6291718 | 5.4862818 | 15.7720618 | 0.0000218 |
| *AAV8-AAV6* | -4.4231862 | -9.5660762 | 0.7197038 | 0.1231536 |
| *AAV9-AAV6* | -1.4798697 | -6.6227597 | 3.6630203 | 0.9621669 |
| *AAV9-AAV8* | 2.9433166 | -2.1995734 | 8.0862066 | 0.5254397 |

**Supplementary Table 7: WT Clone - % GFP-GFAP Tukey multiple comparisons of means**

| Groups compared | Difference between means | Lower bound of confidence interval | Upper bound of confidence interval | adjusted *p*-value |
|---|---|---|---|---|
| AAV1-Control | 6.2260611 | -6.758751 | 19.210873 | 0.708119 |
| *AAV2-Control* | 12.1599764 | -0.8248357 | 25.144789 | 0.0761157 |
| *AAV5-Control* | 1.3743675 | -11.610445 | 14.35918 | 0.9998324 |
| *AAV6-Control* | 7.8391444 | -5.1456677 | 20.823957 | 0.4648422 |
| *AAV8-Control* | 8.1422717 | -4.8425404 | 21.127084 | 0.4215856 |
| *AAV9-Control* | 6.0126894 | -6.9721227 | 18.997501 | 0.7388185 |
| *AAV2-AAV1* | 5.9339153 | -7.0508968 | 18.918727 | 0.7498822 |
| *AAV5-AAV1* | -4.8516936 | -17.836506 | 8.133119 | 0.8807227 |
| *AAV6-AAV1* | 1.6130833 | -11.371729 | 14.597895 | 0.9995776 |
| *AAV8-AAV1* | 1.9162106 | -11.068602 | 14.901023 | 0.9988751 |
| *AAV9-AAV1* | -0.2133717 | -13.198184 | 12.77144 | 1 |
| *AAV5-AAV2* | -10.785609 | -23.770421 | 2.199203 | 0.1469821 |
| *AAV6-AAV2* | -4.320832 | -17.305644 | 8.66398 | 0.9266754 |
| *AAV8-AAV2* | -4.0177047 | -17.002517 | 8.967107 | 0.9470012 |
| *AAV9-AAV2* | -6.147287 | -19.132099 | 6.837525 | 0.7195679 |
| *AAV6-AAV5* | 6.4647769 | -6.5200352 | 19.449589 | 0.6727472 |
| *AAV8-AAV5* | 6.7679042 | -6.2169079 | 19.752716 | 0.6268008 |
| *AAV9-AAV5* | 4.6383219 | -8.3464903 | 17.623134 | 0.9007778 |
| *AAV8-AAV6* | 0.3031273 | -12.681685 | 13.287939 | 1 |
| *AAV9-AAV6* | -1.826455 | -14.811267 | 11.158357 | 0.9991421 |
| *AAV9-AAV8* | -2.1295823 | -15.114395 | 10.85523 | 0.9979697 |

**Supplementary Table 8: AD Clone - % GFP-GFAP Tukey multiple comparisons of means**

| Groups compared | Difference between means | Lower bound of confidence interval | Upper bound of confidence interval | adjusted *p*-value |
|---|---|---|---|---|
| AAV1-Control | 4.981701 | -2.4716342 | 12.4350368 | 0.3500527 |
| *AAV2-Control* | 10.513991 | 3.0606553 | 17.9673262 | 0.0025974 |
| *AAV5-Control* | 2.417285 | -5.0360507 | 9.8706203 | 0.9345128 |
| *AAV6-Control* | 13.240203 | 5.7868674 | 20.6935384 | 0.0001735 |
| *AAV8-Control* | 14.95224 | 7.4989042 | 22.4055752 | 0.0000334 |
| *AAV9-Control* | 6.483686 | -0.9696496 | 13.9370214 | 0.1159479 |
| *AAV2-AAV1* | 5.532289 | -1.921046 | 12.985625 | 0.2417394 |
| *AAV5-AAV1* | -2.564416 | -10.017752 | 4.888919 | 0.9152943 |
| *AAV6-AAV1* | 8.258502 | 0.8051662 | 15.7118371 | 0.0236931 |
| *AAV8-AAV1* | 9.970538 | 2.5172029 | 17.4238739 | 0.0044624 |
| *AAV9-AAV1* | 1.501985 | -5.9513509 | 8.9553201 | 0.9937627 |
| *AAV5-AAV2* | -8.096706 | -15.550041 | -0.6433704 | 0.0276042 |
| *AAV6-AAV2* | 2.726212 | -4.7271233 | 10.1795477 | 0.8905941 |
| *AAV8-AAV2* | 4.438249 | -3.0150865 | 11.8915844 | 0.4805194 |
| *AAV9-AAV2* | -4.030305 | -11.48364 | 3.4230306 | 0.5879697 |
| *AAV6-AAV5* | 10.822918 | 3.3695826 | 18.2762536 | 0.001908 |
| *AAV8-AAV5* | 12.534955 | 5.0816194 | 19.9882904 | 0.0003471 |
| *AAV9-AAV5* | 4.066401 | -3.3869344 | 11.5197366 | 0.5783291 |
| *AAV8-AAV6* | 1.712037 | -5.7412987 | 9.1653722 | 0.9876045 |
| *AAV9-AAV6* | -6.756517 | -14.209853 | 0.6968185 | 0.0922437 |
| *AAV9-AAV8* | -8.468554 | -15.921889 | -1.0152183 | 0.0193973 |

**Supplementary Table 9: Iso Clone - % GFP-GFAP Tukey multiple comparisons of means**

| Groups compared | Difference between means | Lower bound of confidence interval | Upper bound of confidence interval | adjusted *p*-value |
|---|---|---|---|---|
| AAV1-Control | 5.2611503 | -5.9536747 | 16.475975 | 0.7275977 |
| *AAV2-Control* | 10.8794875 | -0.3353376 | 22.094313 | 0.0610605 |
| *AAV5-Control* | 4.6490754 | -6.5657496 | 15.8639 | 0.822052 |
| *AAV6-Control* | 19.3082534 | 8.09342833 | 30.523078 | 0.0002589 |
| *AAV8-Control* | 22.73128 | 11.516455 | 33.946105 | 0.0000288 |
| *AAV9-Control* | 15.8854034 | 4.67057834 | 27.100228 | 0.0024873 |
| *AAV2-AAV1* | 5.6183371 | -5.5964879 | 16.833162 | 0.6666958 |
| *AAV5-AAV1* | -0.6120749 | -11.8269 | 10.60275 | 0.9999966 |
| *AAV6-AAV1* | 14.047103 | 2.83227799 | 25.261928 | 0.0083584 |
| *AAV8-AAV1* | 17.4701297 | 6.25530463 | 28.684955 | 0.0008691 |
| *AAV9-AAV1* | 10.624253 | -0.590572 | 21.839078 | 0.0709217 |
| *AAV5-AAV2* | -6.2304121 | -17.445237 | 4.984413 | 0.5585171 |
| *AAV6-AAV2* | 8.4287659 | -2.7860592 | 19.643591 | 0.2299876 |
| *AAV8-AAV2* | 11.8517925 | 0.63696749 | 23.066618 | 0.0339105 |
| *AAV9-AAV2* | 5.0059159 | -6.2089091 | 16.220741 | 0.7688518 |
| *AAV6-AAV5* | 14.6591779 | 3.4443529 | 25.874003 | 0.0055952 |
| *AAV8-AAV5* | 18.0822046 | 6.86737955 | 29.29703 | 0.0005797 |
| *AAV9-AAV5* | 11.2363279 | 0.02150291 | 22.451153 | 0.0493577 |
| *AAV8-AAV6* | 3.4230266 | -7.7917984 | 14.637852 | 0.9502067 |
| *AAV9-AAV6* | -3.42285 | -14.637675 | 7.791975 | 0.9502184 |
| *AAV9-AAV8* | -6.8458766 | -18.060702 | 4.368948 | 0.4522198 |

### Supplementary Table 10: % GFP Linear mixed effects model relative to AAV6

| AAV Comparison | Estimate | Standard Error | *t*-value | Pr(>\|t\|) | Significance |
|---|---|---|---|---|---|
| *Intercept* | 9.9058 | 1.0439 | 9.489 | 0.00488 | ** |
| *Control* | -9.4267 | 0.5284 | -17.841 | 2.00E-16 | *** |
| *AAV1* | -5.5758 | 0.5284 | -10.553 | 2.00E-16 | *** |
| *AAV2* | -5.7892 | 0.5284 | -10.957 | 2.00E-16 | *** |
| *AAV5* | -8.6217 | 0.5284 | -16.317 | 2.00E-16 | *** |
| *AAV8* | -3.9508 | 0.5284 | -7.477 | 1.17E-10 | *** |
| *AAV9* | -4.2367 | 0.5284 | -8.018 | 1.10E-11 | *** |

Significance codes: 0.001***, 0.01**, 0.05*

### Supplementary Table 11: % GFP-MAP2 Linear mixed effects model relative to AAV6

| AAV Comparison | Estimate | Standard Error | *t*-value | Pr(>\|t\|) | Significance |
|---|---|---|---|---|---|
| Intercept | 10.0946 | 1.294 | 7.801 | 0.00396 | ** |
| Control | -9.9645 | 0.8741 | -11.399 | 2.00E-16 | *** |
| AAV1 | -6.925 | 0.8741 | -7.922 | 1.68E-11 | *** |
| AAV2 | -6.6707 | 0.8741 | -7.631 | 5.98E-11 | *** |
| AAV5 | -9.2244 | 0.8741 | -10.553 | 2.00E-16 | *** |
| AAV8 | -3.3835 | 0.8741 | -3.871 | 0.00023 | *** |
| AAV9 | -2.2475 | 0.8741 | -2.571 | 0.01212 | * |

Significance codes: 0.001***, 0.01**, 0.05*

### Supplementary Table 12: % GFP-GFAP Linear mixed effects model relative to AAV6

| AAV Comparison | Estimate | Standard Error | *t*-value | Pr(>\|t\|) | Significance |
|---|---|---|---|---|---|
| Intercept | 17.154 | 1.682 | 10.198 | 3.05E-08 | *** |
| Control | -13.463 | 2.08 | -6.471 | 8.95E-09 | *** |
| AAV1 | -7.973 | 2.08 | -3.832 | 0.000262 | *** |
| AAV2 | -2.278 | 2.08 | -1.095 | 0.277036 | |
| AAV5 | -10.649 | 2.08 | -5.119 | 2.30E-06 | *** |
| AAV8 | 1.813 | 2.08 | 0.871 | 0.386368 | |
| AAV9 | -4.002 | 2.08 | -1.924 | 0.058204 | |

Significance codes: 0.001***, 0.01**,  0.05*
